# Comparative Computational Docking Analysis of MPP+ and Dopamine Binding to Dopamine Transporter in Human and Zebrafish: Mechanistic Insights into MPTP Neurotoxicity

**DOI:** 10.1101/2025.02.12.637834

**Authors:** Khairiah Razali, Jaya Kumar, Wael M. Y. Mohamed

## Abstract

1-methyl-4-phenyl-1,2,3,6-tetrahydropyridine (MPTP) induces parkinsonism via its toxic metabolite, 1-methyl-4-phenylpyridinium (MPP^+^), which structurally mimics dopamine and enters dopaminergic neurons via the dopamine transporter (DAT). To visually depict this mimicry, we performed ligand-protein docking analysis on two species: humans and zebrafish. Human DAT (PDB ID: Q01959) and zebrafish DAT (UniProt ID: Q90ZV1) proteins were prepared with AutoDock Tools. Meanwhile, MPP+ (PubChem ID: CID_39484) and dopamine (PubChem ID: CID_681) ligands were prepared and optimized using the General AMBER Force Field (GAFF) tool. Docking with AutoDock Vina used identical grid parameters of 40 x 40 x 40 Å, which allows blind docking. Binding affinities and interactions were analyzed and visualized using PyMol and LigPlot+. MPP+ exhibited comparable binding affinities to dopamine in both species (human: -7.2 vs -6.1 kcal/mol; zebrafish: -7.6 vs -6.4 kcal/mol), sharing seven out of nine binding residues in humans and nine out of twelve in zebrafish. Spatial overlap with dopamine binding site suggests MPP+ exploits DAT for neuronal entry. However, static docking limitations warrant dynamic validation. This study provides computational evidence for the dopamine-mimetic mechanism of MPP+ but underscores the need for molecular dynamics (MD) simulation and experimental confirmation.

## Introduction

Parkinson’s disease (PD) is the most prevalent motor neurodegenerative disorder, with its global prevalence increasing by over 120% in the last three decades [1]. By 2040, PD cases are projected to reach 12 million worldwide [2]. Given this rising burden, extensive research has been dedicated to exploring therapeutic strategies to mitigate PD. In preclinical studies, neurotoxin-induced animal models play a crucial role in understanding PD pathogenesis and pathophysiology.

Among these neurotoxins, 1-methyl-4-phenyl-1,2,3,6-tetrahydropyridine (MPTP) was discovered in 1982 as a causative agent of parkinsonism [3]. It induces PD-like symptoms by inhibiting complex I of the mitochondrial electron transport chain (ETC), impairing energy production [4]. MPTP remains one of the most widely used neurotoxins in PD research and is considered a classic model for studying the disease [5]. Once systemically administered, MPTP readily crosses the blood-brain barrier (BBB) and enters the central nervous system (CNS), where it is metabolized into its active neurotoxic form, 1-methyl-4-phenylpyridinium (MPP+). Studies have shown that MPP+ mimics dopamine and enters dopaminergic neurons via the dopamine transporter (DAT) [6, 7].

Zebrafish (*Danio rerio*) has emerged as a valuable model for studying neurodegenerative diseases, including PD, due to its genetic and functional similarities to humans. Additionally, zebrafish offer practical advantages over rodents and primates, such as cost-effectiveness and ease of handling, while also aligning with the 3Rs principles (replacement, reduction, and refinement) [8]. Increasingly, MPTP-induced zebrafish models are being employed in PD research. However, studies on the molecular interactions between MPP^+^ and zebrafish DAT remain limited.

Previous findings, including our own [9, 10], have demonstrated that MPTP induces locomotor deficits and molecular dysfunction in adult zebrafish, supporting its validity as a PD model. Given that MPTP induces parkinsonian symptoms in zebrafish, it is hypothesized that MPP+ interacts with zebrafish DAT in a manner similar to human DAT. In this *in silico* study, molecular docking was performed to computationally evaluate the interactions between MPP+ and DAT in both humans and zebrafish, with comparisons to dopamine-DAT interactions. The study hypothesizes that MPP+ exhibits similar binding affinity and interaction patterns to dopamine in both human and zebrafish DAT, enabling it to mimic dopamine transport mechanisms and exert neurotoxic effects. The findings highlight shared binding properties between MPP^+^ and dopamine in human and zebrafish DAT, further supporting the relevance of zebrafish as a model organism for studying dopaminergic neurotoxicity and PD mechanisms.

## Methods

1-methyl-4-phenylpyridinium (MPP+) is a positively charged metabolite of MPTP and shares structural similarity with the dopamine neurotransmitter, enabling it to enter dopaminergic neurons through DAT protein [11]. We conducted a molecular docking analysis to investigate and visualize the interactions of MPP+ and dopamine with DAT, and to assess the degree of similarity between MPP+ and dopamine in their interactions with DAT. In addition to that, the analysis involved using DAT from two different organisms: humans and zebrafish. Both DAT proteins were employed to study the interactions of MPP+ and dopamine, and to compare the ligand-protein interactions between these two organisms.

### Proteins and Ligands Retrieval

For the proteins, the 3D conformations of human DAT (UniProt ID: Q01959) and zebrafish DAT (UniProt ID: Q90ZV1) were retrieved from the UniProt archive (https://www.uniprot.org/) and downloaded in PDB format. We acknowledge that the existence of zebrafish DAT has only been confirmed at the transcript level, unlike human DAT, which has been validated at the protein level. Nonetheless, the high sequence conservation between zebrafish and human DAT, particularly in key functional domains, supports its structural and functional relevance for comparative molecular docking analysis [12]. Additionally, according to the UniProt database, three computationally mapped potential isoforms of human DAT exist. We selected this specific isoform (UniProt ID: Q01959) because it is designated as the reference proteome and represents the most well-annotated version.

For the ligands, both MPP+ and dopamine 2D structures were acquired from the PubChem database (https://pubchem.ncbi.nlm.nih.gov/) using the ID codes CID_39484 and CID_681, respectively, and downloaded in SDF format (Fig 1). These files were used in the molecular docking analysis to investigate the interactions of MPP+ and dopamine with DAT from both organisms.

**Fig 1.**
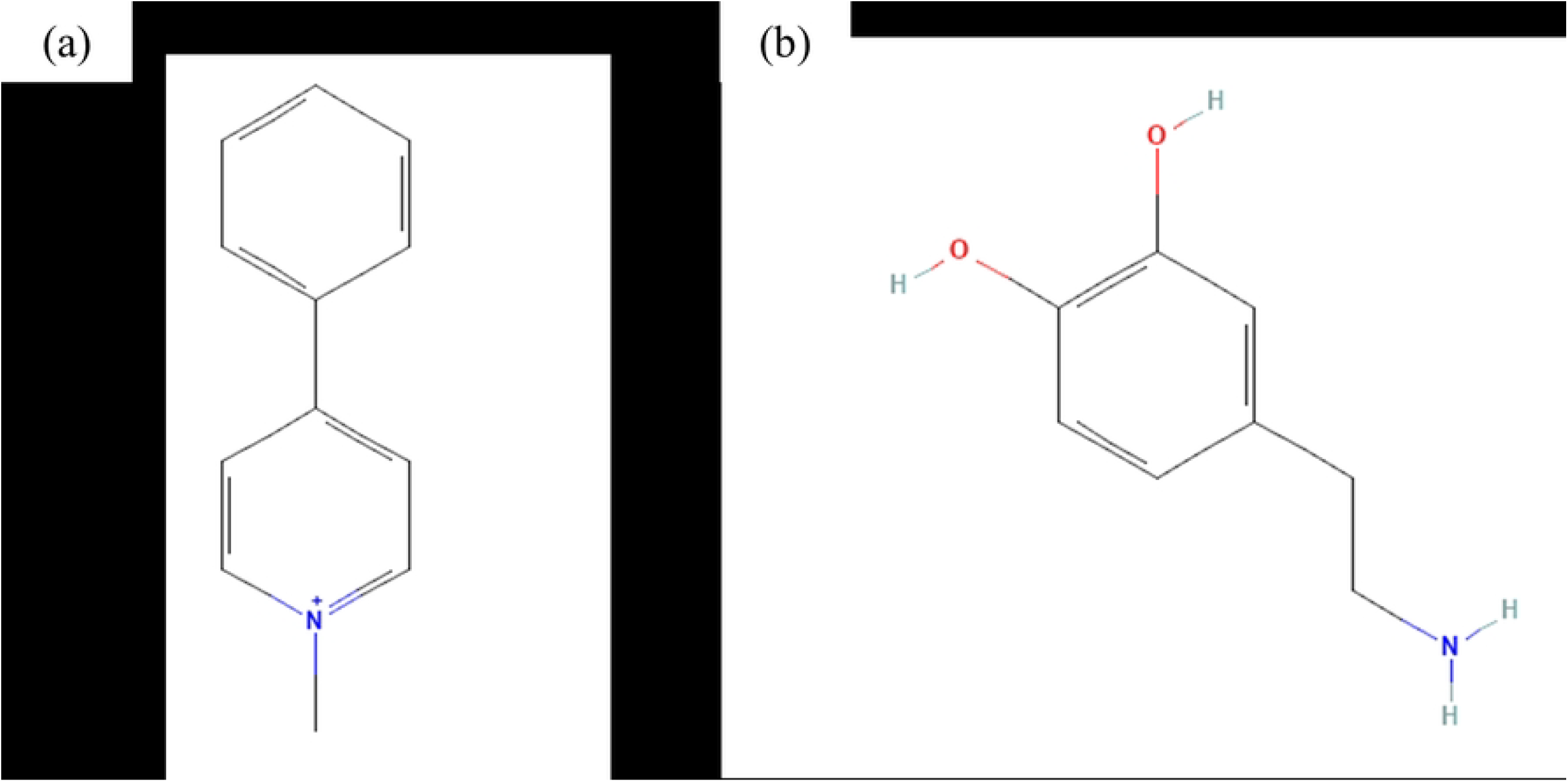
The 2D molecular structures of (a) MPP+ and (b) dopamine retrieved from PubChem database.

### Protein Preparation

The DAT protein from both organisms was prepared for molecular docking using AutoDock Tools software version 1.5.7 (The Scripps Research Institute, CA, USA). Preparation of proteins was carried out by eliminating water molecules and adding up polar hydrogen atoms. Water molecules were removed prior to docking to create a simplified environment that focuses on the direct interactions between the ligand and the binding site of the protein, thus simplifying the docking process and reducing computational complexity [13]. On the other hand, polar hydrogen atoms were added to the proteins to create a more accurate representation of the active site and to facilitate hydrogen bond formations with ligands during the docking process [14].

Therefore, adding polar hydrogen atoms ensures that the protein-ligand complex is properly modelled and improves the reliability of the docking predictions. To further improve the accuracy and reliability of docking process, the atomic interactions within the proteins were parametrized by adding Kollman charges. Prepared proteins were saved in PDBQT format.

### Ligand Preparation

Since dopamine and MPP+ were downloaded in 2D conformation and SDF format, it needed an additional preparation process. Firstly, the 2D structure was converted to 3D conformation using Avogadro software version 1.99.0 [15]. The 3D conformation was optimized utilizing steepest descent calculations with the General AMBER Force Field (GAFF) in the auto-optimization tool. GAFF is a force field option widely used to optimize drug geometries [15]. The optimized 3D conformation of dopamine and MPP+ was saved in PDB format.

After optimization, both ligands underwent ligand preparation using AutoDock Tools software by adding Gasteiger charges and allowing torsions to rotate. Gasteiger charges are commonly used to assign partial atomic charges to ligands or small molecules [16]. The prepared ligands were saved in PDBQT format.

It is essential to highlight that MPP^+^ remains a positively charged cation due to its pyridinium structure [17]. While the addition of Gasteiger charges assigns partial atomic charges based on electronegativity equilibration [18], this process does not modify the fundamental protonation state of MPP+. As a result, MPP^+^ retains its positive charge throughout ligand preparation and docking, ensuring accurate electrostatic interactions with DAT proteins.

### Protein-Ligand Docking and Analysis

Both protein and ligand in PDBQT format were loaded to AutoDock Tools to obtain the grid box parameter. This parameter is important as it defines the size and position of the 3D grid within the protein that is needed for the docking process. In this study, the grid box size was set to the default 40 × 40 × 40 Å, enabling blind docking. This approach allows the ligands to freely explore potential interactions across the protein without being restricted to a predefined binding site. The grid box position for DAT of each organism is stated in Table 1.

**Table 1.** Grid box coordinates for DAT protein of each organism.

| Protein (*Organism*) | UniProt ID | Coordinate x, y, z |
| --- | --- | --- |
| DAT (*Homo sapiens*) | Q01959 | -42.465, -0.569, 55.058 |
| DAT (*Danio rerio*) | Q90ZV1 | -5.125, 1.99, 4.587 |

The docking jobs were conducted using AutoDock Vina version 1.2.0 (The Scripps Research Institute, CA, USA) by executing specific command lines. A single docking run was performed, generating nine potential binding poses (modes). The mode with the lowest binding energy and RMSD (Root Mean Square Deviation) values were selected as the final result as it represents the most thermodynamically favorable and stable ligand-protein interaction, indicating a higher likelihood of biological relevance.

For the result analysis and visualization, the original protein PDB file and the PDBQT output file were superimposed using PyMOL software version 2.5.5 (The PyMOL Molecular Graphics System, Schrödinger, Inc) to generate a 3D diagram of the ligand-protein complex. The ligand sites preset was chosen to highlight the interaction sites between the ligand and the protein, and the resulting complex was saved in PDB format. Subsequently, the saved complex was further analyzed using LigPlot+ software version 2.2.8 [19], to generate a 2D diagram depicting the ligand-protein interactions in detail.

## Results

1-methyl-4-phenylpyridinium (MPP+) bears a structural resemblance to the dopamine neurotransmitter, allowing it to undergo transportation into dopaminergic neurons through the DAT protein. To illustrate this similarity in their interaction with the DAT protein, we conducted a ligand-protein docking analysis. Table 2 provides comprehensive information regarding the binding of dopamine and MPP+ to the DAT protein in both human and zebrafish, while Figs 2 to 5 provide visual depictions of these binding interactions on the DAT protein for the respective species.

**Table 2.** Molecular interaction results of DAT protein with the active ligands dopamine and MPP+.

| Organisms / Parameters | *Homo sapiens* (UniProt ID: Q01959) |  | *Danio rerio* (UniProt ID: Q90ZV1) |  |
| --- | --- | --- | --- | --- |
|  | Dopamine | MPP+ | Dopamine | MPP+ |
| Binding affinity (kcal/mol) | -6.5 | -7.6 | -6.4 | -7.6 |
| H-bond interacting residues | **ASP79** | - | **ASP95**<br>SER165 | - |
| Hydrophobic interacting residues | SER149<br>**VAL152**<br>GLY153<br>**TYR156**<br>**PHE326**<br>SER422<br>ALA423<br>GLY426 | PHE76<br>**ASP79**<br>ALA81<br>ARG85<br>**VAL152**<br>**TYR156**<br>PHE320<br>**PHE326**<br>ASP476 | **PHE92**<br>**VAL168**<br>**GLY169**<br>**TYR172**<br>**PHE335**<br>**SER431**<br>**ALA432**<br>**GLY435** | **PHE92**<br>**ASP95**<br>**VAL168**<br>**GLY169**<br>**TYR172**<br>PHE329<br>**PHE335**<br>**SER431**<br>**ALA432**<br>**GLY435**<br>MET436 |

\*Dopamine PubChem ID: 681; MPP+ PubChem ID: 39484
\*Same interacting residues are in bold.

**Fig 2.**
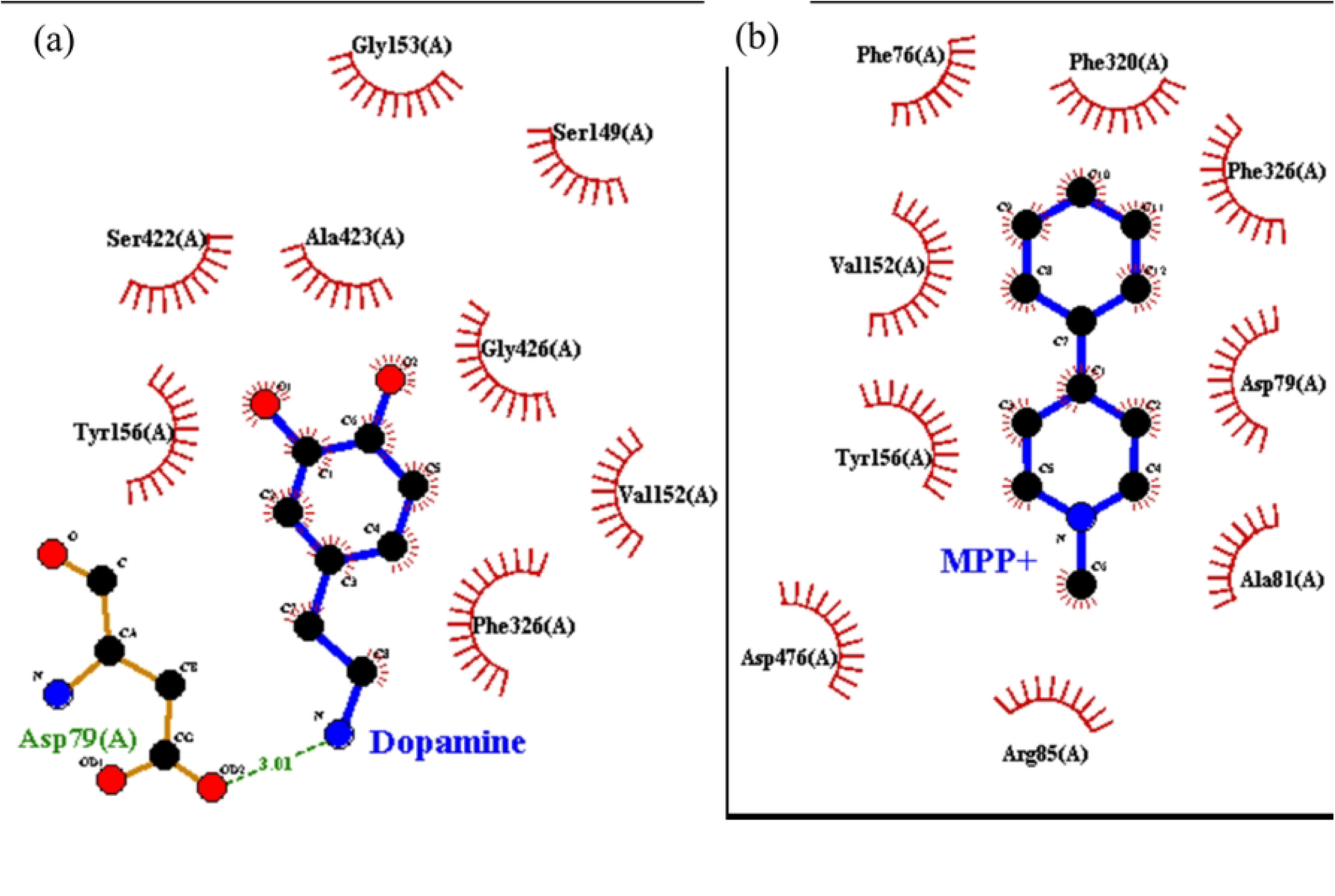
Molecular docking of human DAT protein with dopamine and MPP+ ligands. The 2D diagrams show the binding between DAT and a) dopamine and b) MPP+. Blue stick models indicate the ligand, brown stick models indicate important residues near the binding site, green dotted lines indicate the hydrogen bonds, and half-moon dashes indicate the hydrophobic contacts between residue and ligand.

Dopamine side chains formed one hydrogen and eight hydrophobic interactions with the human DAT protein. Specifically, the C8 nitrile group established a hydrogen bond with Asp79 residue. Additionally, the dopamine side chains engaged in hydrophobic interactions with the Ser149, Val152, Gly153, Tyr156, Phe326, Ser422, Ala423, and Gly426 residues, as illustrated in Fig 2a. For MPP+, the methyl, phenyl, and pyridinium groups engaged in nine hydrophobic interactions with Phe76, Asp79, Ala81, Arg85, Val152, Tyr156, Phe320, Phe326, and Asp476 residues in the human DAT protein (Fig 2b). Markedly, dopamine and MPP+ share a commonality in binding to four out of nine of the same residues in the DAT interaction as indicated in green in Table 2.

Both dopamine and MPP+ demonstrate comparable affinity for binding to the DAT protein, with energies of -6.5 and -7.6 kcal/mol, respectively. Most importantly, their binding occurs at the same spatial location (Fig 3), providing further evidence of their remarkably similar interactions with the amino acid residues of the protein.

**Fig 3.**
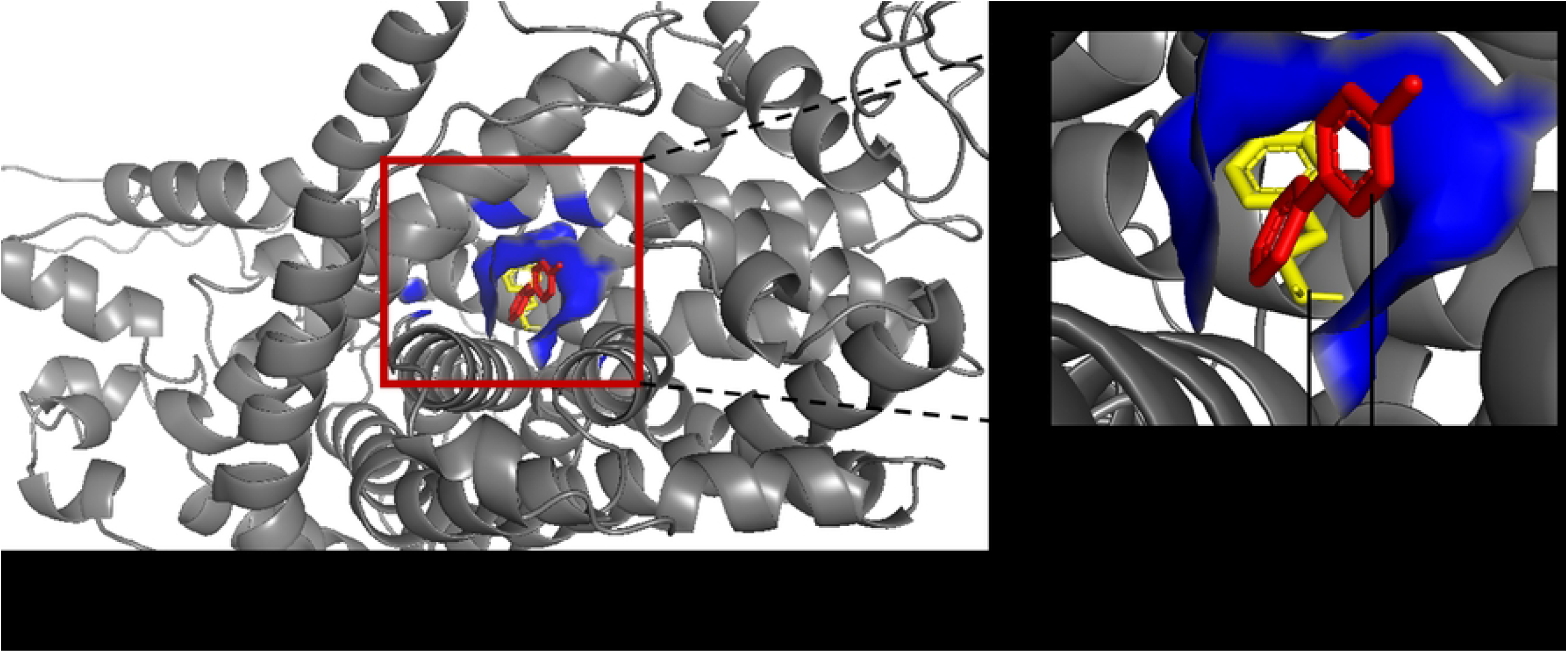
The superimposed 3D diagram shows the binding sites of dopamine and MPP+ on human DAT protein. The grey ribbon structure represents the DAT protein, and the blue solid structure highlights the location of the binding sites. The yellow and red stick models represent dopamine and MPP+, respectively.

Similar to its interaction with the human DAT protein, MPP+ side chains also formed hydrophobic contacts with the zebrafish DAT protein. Specifically, it interacted with eleven residues: Phe92, Asp95, Val168, Gly169, Tyr172, Phe329, Phe335, Ser431, Ala432, Gly435, and Met436 (Fig 4b). It is important to mention that these nine residues are also present in the dopamine-DAT complex (Table 2). Dopamine established two hydrogen bonds with the DAT protein, one link at the C8 nitrile group and another at the C6 carbonyl group, and eight hydrophobic bonds with Phe42, Val168, Gly169, Tyr172, Phe335, Ser431, Ala432, and Gly435 (Fig 4a).

**Fig 4.**
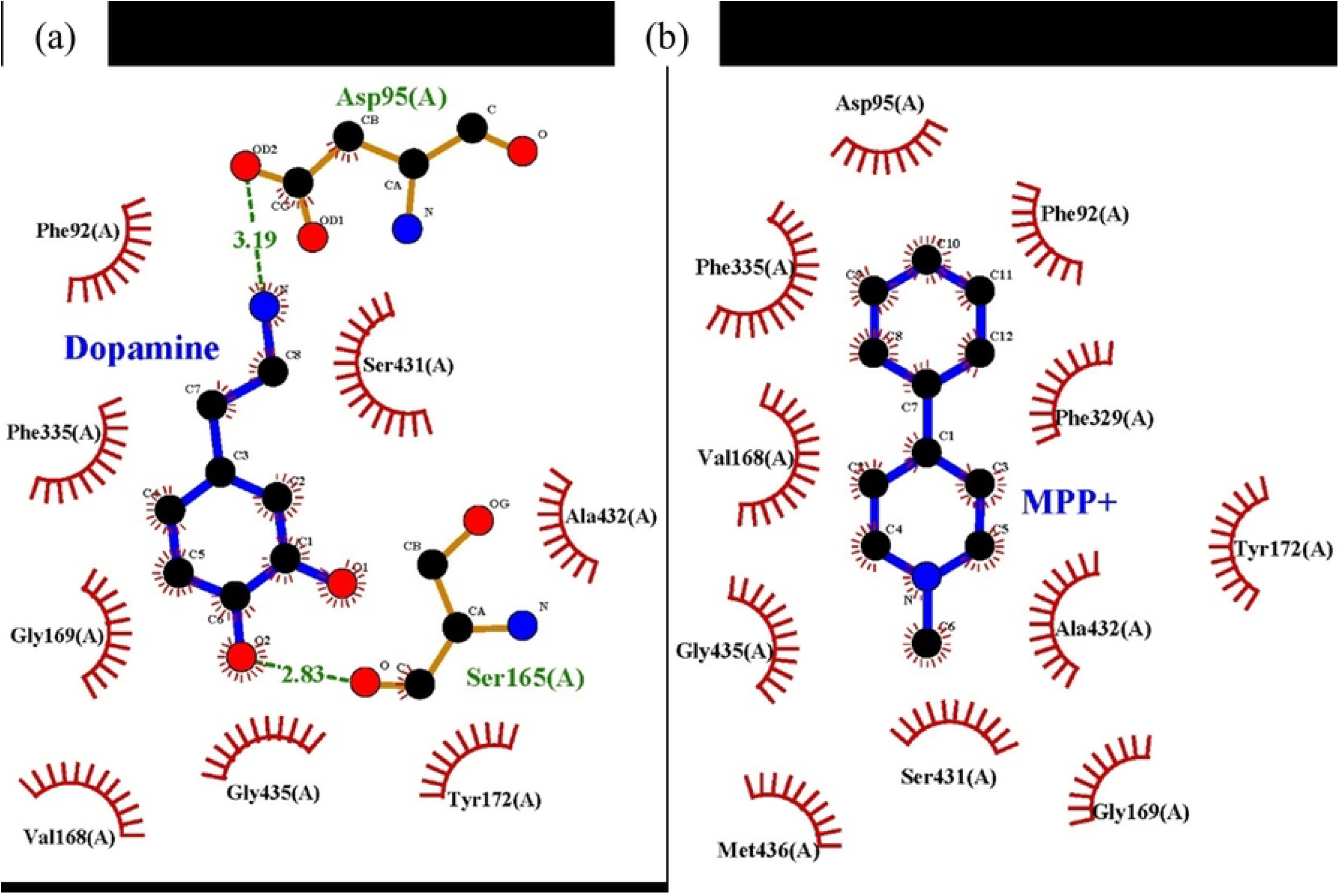
Molecular docking of zebrafish DAT protein with dopamine and MPP+ ligands. The 2D diagrams show the binding between DAT and a) dopamine and b) MPP+. Blue stick models indicate the ligand, brown stick models indicate important residues near the binding site, green dotted lines indicate the hydrogen bonds, and half-moon dashes indicate the hydrophobic contacts between residue and ligand.

The binding affinity of the MPP+-DAT complex closely resembles that of the dopamine-DAT complex, with values of -7.6 kcal/mol and -6.4 kcal/mol, respectively. In terms of spatial location, MPP+ binds to the zebrafish DAT protein at a location strikingly similar to dopamine (Fig 5).

**Fig 5.**
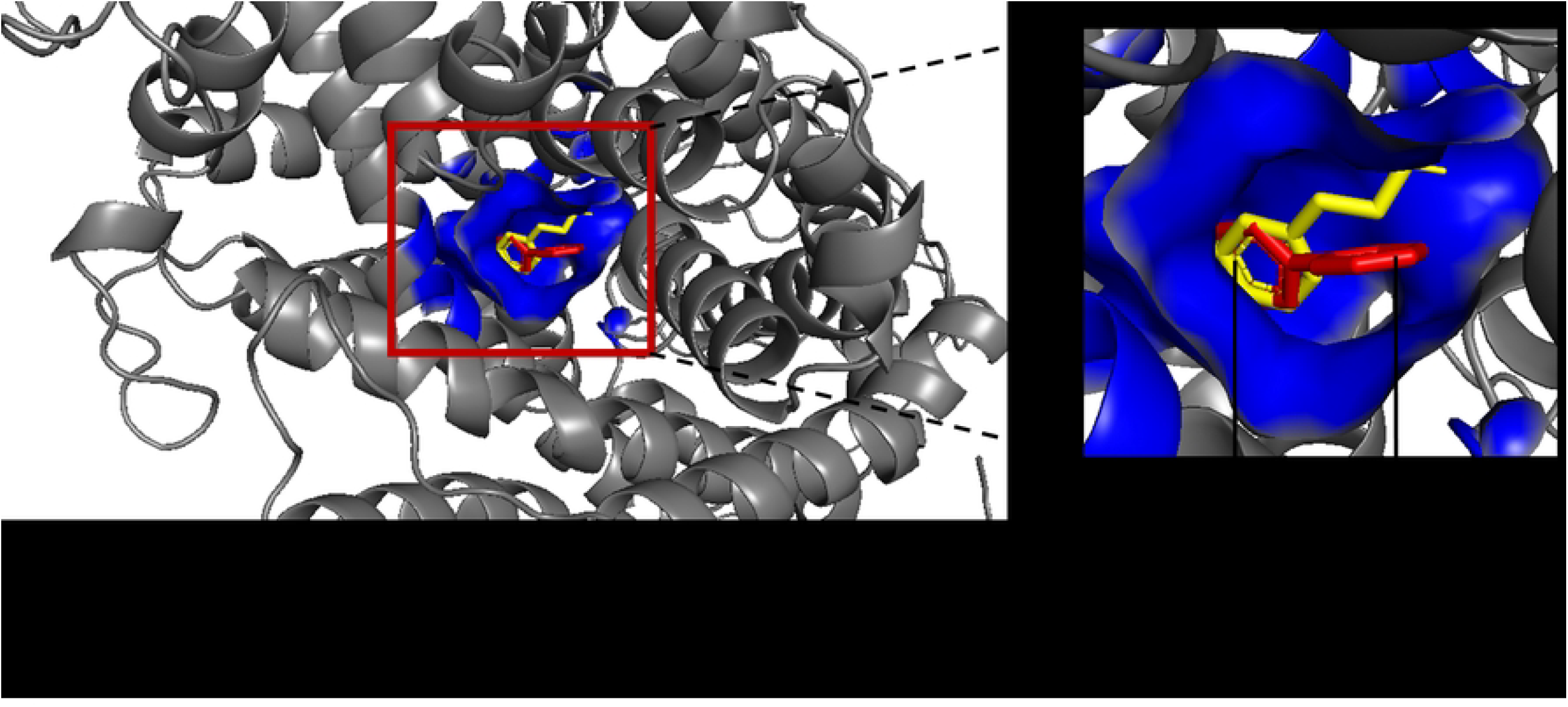
The superimposed 3D diagram shows the binding sites of dopamine and MPP+ on zebrafish DAT protein. The grey ribbon structure represents the DAT protein, and the blue solid structure highlights the location of the binding sites. The yellow and red stick models represent dopamine and MPP+, respectively.

## Discussion

We conducted molecular docking analysis on the neurotoxic metabolite, MPP+, to elucidate its interaction with the DAT protein. Subsequently, we conducted a comparison between this particular interaction and the interaction between dopamine and DAT to determine if MPP+ interacts with the same residues of DAT as dopamine. Through our investigation, we discovered that MPP+ interacts with similar protein residues of the human DAT protein that dopamine does while attempting to enter the dopaminergic neurons. Moreover, MPP+ exhibits a slightly greater affinity for binding DAT protein relative to dopamine. A parallel observation was found in the zebrafish DAT protein. These findings offer more evidence that MPP+ imitates the process by which dopamine binds, enabling it to efficiently manipulate of the dopamine uptake system and enter dopaminergic neurons. In addition, the discovery of similarities between both species provides evidence that the function of the DAT protein is preserved across different species. This suggests that zebrafish is a suitable model organism for studying the neurotoxic effects of MPTP.

MPP+ is a metabolically active product that is produced when MPTP is broken down by monoamine oxidase B (MAO-B) in glial cells. The compound is an organic molecule that is positively charged, and its molecular formula is C12H12N+. The chemical composition consists of an aromatic ring, specifically a benzene ring, connected to a six-membered ring comprising of a nitrogen atom and five carbon atoms, commonly referred to as a pyridinium ring [17] (Fig 1a). Meanwhile, dopamine is a catecholamine with a molecular formula C8H11NO2. Like MPP+, dopamine also has a benzene ring, but instead of attached to a pyridinium ring, its benzene ring possesses hydroxyl (-OH) groups [20] (Fig 1b). MPP+ possesses a pyridinium ion (-N+) with a positive charge, whereas dopamine contains an amine group (-NH2) that can be protonated to generate a positively charged amine (-NH3+). While the structure of MPP+ is not exactly the same as dopamine, it possesses similar important characteristics such as a benzene ring and a positive charge. These qualities enable MPP+ to engage with the same binding site on the DAT protein, thereby imitating dopamine entry into dopaminergic neurons. The interaction data from our study offers a comprehensive analysis of the specific DAT residues that play a role in binding dopamine and MPP+. This comprehensive mapping will assist researchers in gaining a deeper understanding of the molecular insights into MPTP-induced neurotoxicity.

The DAT protein is a transmembrane protein that regulates dopamine levels by controlling the uptake of extracellular dopamine and terminating dopaminergic transmission. MPP+ induces the translocation of the DAT protein receptor to the cell surface in dopaminergic neurons, resulting in an increase in MPP+ uptake [13]. This clarifies the precise toxicity of MPP+ towards dopaminergic neurons. MPP+ induces neurotoxicity by blocking complex I of the mitochondrial ETC in neurons, hence altering oxidative phosphorylation in mitochondria. The disruption of the mitochondrial ETC is detrimental since it leads to a decrease in adenosine triphosphate (ATP) levels. In addition, MPP+ disturbs the equilibrium of dopamine in dopaminergic neurons by affecting both its concentration and release [11].

Another significant discovery was the parallel detection of the interactions between dopamine and MPP+ with the DAT protein in both human and zebrafish. Additionally, we discovered that MPP+ interacts with the identical amino acid residues as dopamine, not only in the human DAT protein but also in the zebrafish DAT protein. The preservation of DAT protein function across different species indicates its crucial and essential role in the process of dopamine neurotransmission. In addition, it is important to note that the architecture and spatial binding sites differ between the human and zebrafish DAT protein (Figs 3 and 5). However, they both share the same function of binding MPP+ and dopamine at the same interaction residues. This implies that evolutionary conservation serves a functional purpose rather than a structural one. This finding aligns with a prior investigation conducted by Shumay, Fowler and Volkow (21), which revealed that the DAT exhibits a high degree of evolutionary conservation at the protein level but is less conserved at the genome level.

The fact that MPP+ and dopamine bind to the zebrafish DAT protein in a comparable manner as they do to the human DAT protein suggests that the zebrafish is a good model organism for studying dopaminergic neurotoxicity and the mechanisms of PD. This resemblance suggests that the discoveries about dopamine uptake mechanism in zebrafish are likely to have relevance and applicability to human biology. By leveraging the regenerative capacity of the zebrafish to repair cells in the CNS [22], we can utilize this understanding to investigate methods for reversing dopaminergic neurodegeneration in PD. Given that PD is a degenerative disease, it is worth exploring the possibility of regenerating dopaminergic neurons in zebrafish in order to restore dopamine neurotransmission. The results of such studies may have potential implications for developing regenerative therapies for PD in humans, offering novel approaches for treatment that could mitigate the progression of the disease.

Collectively, the molecular docking analysis conducted on both human and zebrafish DAT proteins offers convincing evidence that MPP+ enters dopaminergic neurons by binding to DAT protein residues that closely resemble those utilized by dopamine. These findings provide further support for the hypotheses suggesting that the neurotoxin MPTP exerts its toxic effects on dopaminergic neurons via its metabolite MPP+. This occurs through the mimicry of dopamine neurotransmitter structure, enabling MPP+ to bind to the DAT protein and access the neurons.

## Limitations and Conclusion

While our findings have effectively shown the commonalities in the interaction between MPP+ and dopamine with DAT protein residues, the analysis has limitations in terms of its depth. *In silico* findings are limited to predicting the interaction and do not consider the intricacies of physiological systems. Similar binding site occupancy suggests a structural similarity in MPP+ and dopamine interactions with DAT but does not necessarily confirm functional mimicry. Similarly, binding affinity values offer theoretical estimates of the interaction strength between MPP+ and dopamine with DAT, however, these values are not inherently biologically meaningful on their own unless integrated with other computational and experimental approaches. To obtain more definitive interpretations, it is necessary to conduct research both *in vitro* and *in vivo* to compare the interactions of dopamine-DAT and MPP+-DAT in a realistic physiological setting. Multiple studies have utilized MPP+ to investigate DAT activity [23, 24]. Nevertheless, there remains a scarcity of research that directly and exclusively examines the binding affinity and competitive nature of dopamine and MPP+ towards the DAT protein, especially in zebrafish models.

Additionally, our analysis did not account for the temporal dynamics of binding. Our understanding is limited to the binding affinity at a specific moment and within a static conformation. The duration of intracellular transport for MPP+ after binding to the DAT protein, as well as how this process mechanistically compares to dopamine transport, remains uncertain. To overcome this limitation, MD simulations and laboratory experiments are needed to provide deeper insights into stability, conformational changes, and transport mechanisms of these interactions over time.

Lastly, while the docking results suggest that MPP+ binds similarly to dopamine in both species, our analysis remains limited by the static nature of molecular docking. Docking primarily provides a snapshot of ligand-protein interactions at a fixed conformation and does not account for protein flexibility, solvent effects, or dynamic binding processes that occur in a cellular environment. Additionally, the accuracy of docking results depends on the quality of the input protein structure, which in the case of zebrafish DAT, has only been validated at the transcript level rather than at the protein level. Integrating sophisticated AI-based analyses could significantly improve data interpretation and overcome some of these challenges. For instance, AI-driven structure prediction tools such as AlphaFold could enhance the accuracy of zebrafish DAT structural modeling by predicting its three-dimensional conformation with high confidence, even in the absence of experimental protein validation. Moreover, deep learning algorithms, particularly graph neural networks (GNNs)—could refine docking predictions by learning from vast datasets of molecular interactions. GNNs can capture complex spatial relationships between ligands and proteins, improving binding pose predictions and affinity estimations.

Our study demonstrates that MPP+ binds to the DAT protein in both species at a location similar to that of dopamine. This molecular docking analysis of human and zebrafish DAT proteins provides predictive evidence that the neurotoxin MPTP exerts its toxic effects on dopaminergic neurons through its metabolite MPP+, which mimics the structure of dopamine, allowing it to bind to DAT and enter the neurons. Demonstrating that MPP^+^ and dopamine bind to zebrafish DAT in a similar manner to human DAT supports the validity of zebrafish as a model organism for studying dopaminergic neurotoxicity and PD mechanisms. This similarity implies that findings in dopamine uptake mechanism in zebrafish are likely to be relevant and translatable to human system. With the zebrafish regenerative ability to regenerate cells in the CNS, we can exploit this knowledge to study how to reverse dopaminergic neurodegeneration in PD. Since PD is a progressive neurodegenerative disease, we can try to regenerate dopaminergic neurons in zebrafish to salvage dopamine neurotransmission, and the findings may not be impossible to translate to human biology. Our future work will integrate docking with MD simulations, enhanced by AI-assisted trajectory analysis and validated through experimental studies, to achieve a more comprehensive evaluation of MPP+/DAT and dopamine/DAT binding stability over time, providing a more accurate representation of real biological conditions.

## Supporting Information

## References

1. Pereira GM, Teixeira-dos-Santos D, Soares NM, Marconi GA, Friedrich DC, Saffie Awad P, et al. A systematic review and meta-analysis of the prevalence of Parkinson’s disease in lower to upper-middle-income countries. npj Parkinson’s Disease. 2024;10(1):181.

2. Dorsey ER, Sherer T, Okun MS, Bloem BR. The emerging evidence of the Parkinson pandemic. Journal of Parkinson’s disease. 2018;8(S1):S3-S8.

3. Langston JW. The MPTP story. Journal of Parkinson’s disease. 2017;7(1):S11–S9.

4. Ferrucci M, Fornai F. MPTP neurotoxicity: actions, mechanisms, and animal modeling of Parkinson’s disease. Handbook of Neurotoxicity: Springer; 2021. p. 1–41.

5. AlShimemeri S, Di Luca DG, Fox SH. MPTP Parkinsonism and implications for understanding Parkinson’s disease. Movement Disorders Clinical Practice. 2021;9(1):42.

6. Waerzeggers Y, Monfared P, Viel T, Winkeler A, Jacobs AH. Mouse models in neurological disorders: applications of non-invasive imaging. Biochimica et Biophysica Acta (BBA)-Molecular Basis of Disease. 2010;1802(10):819–39.

7. Wen S, Aki T, Unuma K, Uemura K. Chemically induced models of Parkinson’s disease: history and perspectives for the involvement of ferroptosis. Frontiers in cellular neuroscience. 2020;14:581191.

8. Lauwereyns J, Bajramovic J, Bert B, Camenzind S, De Kock J, Elezović A, et al. Toward a common interpretation of the 3Rs principles in animal research. Lab Animal. 2024:1–4.

9. Razali K, Mohd Nasir MH, Kumar J, Mohamed WM. Mitophagy: A Bridge Linking HMGB1 and Parkinson’s Disease Using Adult Zebrafish as a Model Organism. Brain Sciences. 2023;13(7):1076.

10. Razali K, Mohd Nasir MH, Othman N, Doolaanea AA, Kumar J, Nabeel Ibrahim W, Mohamed WM. Characterization of neurobehavioral pattern in a zebrafish 1-methyl-4-phenyl-1, 2, 3, 6-tetrahydropyridine (MPTP)-induced model: A 96-hour behavioral study. PLoS One. 2022;17(10):e0274844.

11. Choi SJ, Panhelainen A, Schmitz Y, Larsen KE, Kanter E, Wu M, et al. Changes in neuronal dopamine homeostasis following 1-methyl-4-phenylpyridinium (MPP+) exposure. Journal of Biological Chemistry. 2015;290(11):6799–809.

12. Wang Y, Takai R, Yoshioka H, Shirabe K. Characterization and expression of serotonin transporter genes in zebrafish. The Tohoku journal of experimental medicine. 2006;208(3):267–74.

13. Agu P, Afiukwa C, Orji O, Ezeh E, Ofoke I, Ogbu C, et al. Molecular docking as a tool for the discovery of molecular targets of nutraceuticals in diseases management. Scientific Reports. 2023;13(1):13398.

14. Azad I, Khan T, Maurya AK, Irfan Azad M, Mishra N, Alanazi AM. Identification of severe acute respiratory syndrome coronavirus-2 inhibitors through in silico structure-based virtual screening and molecular interaction studies. Journal of Molecular Recognition. 2021;34(10):e2918.

15. Hanwell MD, Curtis DE, Lonie DC, Vandermeersch T, Zurek E, Hutchison GR. Avogadro: an advanced semantic chemical editor, visualization, and analysis platform. Journal of cheminformatics. 2012;4:1–17.

16. Che X, Liu Q, Zhang L. An accurate and universal protein-small molecule batch docking solution using Autodock Vina. Results in Engineering. 2023;19:101335.

17. Lehner A, Johnson M, Simkins T, Janis K, Lookingland K, Goudreau J, Rumbeiha W. Liquid chromatographic–electrospray mass spectrometric determination of 1-methyl-4-phenylpyridine (MPP+) in discrete regions of murine brain. Toxicology Mechanisms and Methods. 2011;21(3):171–82.

18. Oliferenko AA, Pisarev SA, Palyulin VA, Zefirov NS. Atomic charges via electronegativity equalization: generalizations and perspectives. Advances in Quantum Chemistry. 2006;51:139–56.

19. Wallace AC, Laskowski RA, Thornton JM. LIGPLOT: a program to generate schematic diagrams of protein-ligand interactions. Protein engineering, design and selection. 1995;8(2):127–34.

20. Faleye OS, Boya BR, Lee J-H, Choi I, Lee J, Page C. Halogenated antimicrobial agents to combat drug-resistant pathogens. Pharmacological Reviews. 2024;76(1):90–141.

21. Shumay E, Fowler JS, Volkow ND. Genomic features of the human dopamine transporter gene and its potential epigenetic states: implications for phenotypic diversity. PloS one. 2010;5(6):e11067.

22. Godoy R, Hua K, Kalyn M, Cusson V-M, Anisman H, Ekker M. Dopaminergic neurons regenerate following chemogenetic ablation in the olfactory bulb of adult Zebrafish (Danio rerio). Scientific reports. 2020;10(1):12825.

23. Luk B, Mohammed M, Liu F, Lee FJ. A physical interaction between the dopamine transporter and DJ-1 facilitates increased dopamine reuptake. PLoS One. 2015;10(8):e0136641.

24. Sun Y, Selvaraj S, Pandey S, Humphrey KM, Foster JD, Wu M, et al. MPP+ decreases store-operated calcium entry and TRPC1 expression in Mesenchymal Stem Cell derived dopaminergic neurons. Scientific reports. 2018;8(1):11715.

